# Proximity-based super-resolution imaging enabled by DNA base-stacking interactions

**DOI:** 10.1101/2025.06.14.659733

**Authors:** Abhinav Banerjee, Saanya Yadav, Vedanth Shree Vidwath, Simanta Kalita, Sarit S. Agasti, Himanshu Joshi, Mahipal Ganji

**Affiliations:** Department of Biochemistry, Indian Institute of Science, Malleswaram, Bangalore-560012, India; Department of Biotechnology, IIT Hyderabad, Kandi, Telangana 502284, India; New Chemistry Unit and Chemistry and Physics of Materials Unit, Jawaharlal Nehru Centre for Advanced Scientific Research, Bengaluru, India

**Author notes:** Equal contribution (Abhinav Banerjee, Saanya Yadav, Vedanth Shree Vidwath).

**Keywords:** DNA base-stacking, DNA-PAINT, Super-resolution imaging, Molecular dynamics, DNA Origami, Kinetic analysis, Fluorescence

## Abstract

Super-resolution imaging has gained significant traction in recent years due to its unprecedented ability to visualize target biomolecules at nanometer resolution. Here, we demonstrate the capability to detect target pairs that are in close proximity by exploiting base-stacking interactions between two DNA strands, each labelling one target. Our DNA probes hybridize transiently with the each other only when both target molecules are proximal, thus creating a hybridization site for fluorophore-conjugated DNA strand, called imager. In this design, hybridization and co-axial base-stacking act synergistically to enable imager binding, with stacking interactions providing essential stabilization that allows for transient hybridization. This synergy generates the stochastic binding events required for DNA-PAINT imaging, which we call Stack-Proximity-PAINT (Stack-pPAINT). To gain mechanistic insights into hybridization of our probe, we performed atomistic equilibrium and steered molecular dynamics (MD) simulations. The simulations reveal that DNA base-stacking and fluorophore stacking together stabilize the imager. We utilized programmable DNA nanostructures to benchmark the applicability of Stack-pPAINT. As a cellular proof of concept, we visualized microtubular structures using Stack-pPAINT with antibodies targeting both alpha- and beta-tubulin molecules. This probe technology offers promising applications in cell biology research aimed at elucidating spatial interactomes at high resolution within cells.

## Introduction

Every single functionality within a cell, be it prokaryotes or eukaryotes, is orchestrated by many protein molecules that closely interact with each other either directly or indirectly. These interactions occur at proximity to one another within the cell, thus enabling the intricate control over cellular processes. Detecting these interactions at greater spatiotemporal resolution has been a field that is actively evolving. Co-immunoprecipitation (Co-IP), yeast-two-hybrid, protein affinity chromatography and other such biochemical techniques have helped reveal great details on protein-protein interactions.^1^ These techniques excel in even detecting weaker interactions but fall short when one needs to understand the place within the cell where these interactions occur. With the advent of microscopy, FRET (Förster resonance energy transfer) based techniques allow for the detection of these interactions if the partners are present within a few nanometers (Förster Radii) from each other. This technique strongly relies on the ability to generate fluorescently labelled proteins or use antibodies with specific FRET pairs that can then be used to stain these proteins in fixed cells. This technique provides diffraction limited information for the exact position of the interacting pairs. Similarly, PLA (Proximity Ligation Assay) relies on DNA conjugated antibodies. These DNA strands get ligated with each other, enzymatically amplified, and would thus give signal if they are present within 30 – 40 nm from each other. This also allows for diffraction limited location information for the presence of protein interacting pairs. This low-resolution information, though valuable, keeps us in the dark regarding the density of these partners, separation between them and the finer distribution of these pairs within the cell.

Super-resolved microscopic tools to discern such partners at greater resolution would circumvent these drawbacks of FRET and PLA. Previous approaches have relied on exploiting DNA-PAINT (Points Accumulation for Imaging in Nanoscale Topography)^2^ with split docking strands, where the imager molecule binding is facilitated only when both parts of the docking strands hybridize via their stem portions to allow for a T-shaped platform for the imager to bind^3^, or a blocked docking site which is exposed by strand invasion by a nearby partner DNA^4^. These approaches are reliable strategies but bare unpredictable kinetics due to the role of multiple DNA strands that interact in a non-canonical manner.

In this study we outline a super-resolution imaging approach to detect protein-protein interaction pairs with the help of DNA-PAINT aided by stacking interactions to tune the detection of binding events. Conventionally, DNA-PAINT relies on the transient hybridization of DNA imager molecule with its complimentary docking strand for time durations that are detectable by a chip-based camera. We exploit the benefits provided by stacking interactions across a nick site adjacent to the site of DNA binding, that stabilizes the binding of imager molecules^5^. In the absence of this stacking interaction, the kinetics of this molecule is tuned such that there is no detectable signal upon binding between the docking strand and the imager molecule.

We have tuned the binding times such that only in the presence of the stacking pair, the binding event is stabilized sufficiently for it to be detected. To further explain the subtle but profound interactions occurring at the molecular level, we created all-atom models closely mimicking the experimental constructs and performed equilibrium and advanced sampling molecular dynamics (MD) simulations. Several microseconds long MD simulations enable us to visualize the nanoscale structures, dynamics and interactions of proximity-PAINT set up. As our approach takes advantage of stacking interactions to attain DNA-PAINT imaging of proximal molecules, we call it Stack-pPAINT (Stacking enabled Proximity-DNA-PAINT). We validate the design of Stack-pPAINT using designer DNA nanostructures and then showcase their cellular application on the microtubular network composed of α-tubulin and β-tubulin heterodimers.

## Results

### Design principle of stacking-assisted proximity detection using DNA-PAINT imaging

Regular DNA-PAINT super-resolution imaging is carried out by acquisition of the transient binding events of fluorophore-attached DNA (imager) strand on its complementary (docking) strand for generating the required “blinking” events for super-resolved reconstruction of the data.^2^ Our recent study showed that an additional dinucleotide base-stacking interaction can significantly stabilize the imager strand (5- to 250-fold) on its docking strand in DNA-PAINT super-resolution imaging.^5^ In this report, we intended to design a super-resolution imaging sensor that specifically results in a detectable signal only when both docking strand and additional stacking interaction are present. To achieve this, we designed a six nucleotide (nt) docking strand which is too short to show any detectable imager binding. However, in the presence of an additional dinucleotide base-stacking interaction at the imager’s terminus via a stem DNA strand, we have fine-tuned interacting DNA strands that facilitate imager binding signal. As the detection is facilitated by DNA hybridization and base-stacking interactions, we named it Stacking-assisted Proximity-detection by DNA-PAINT (Stack-pPAINT) (Figure 1a).

**Figure 1:**
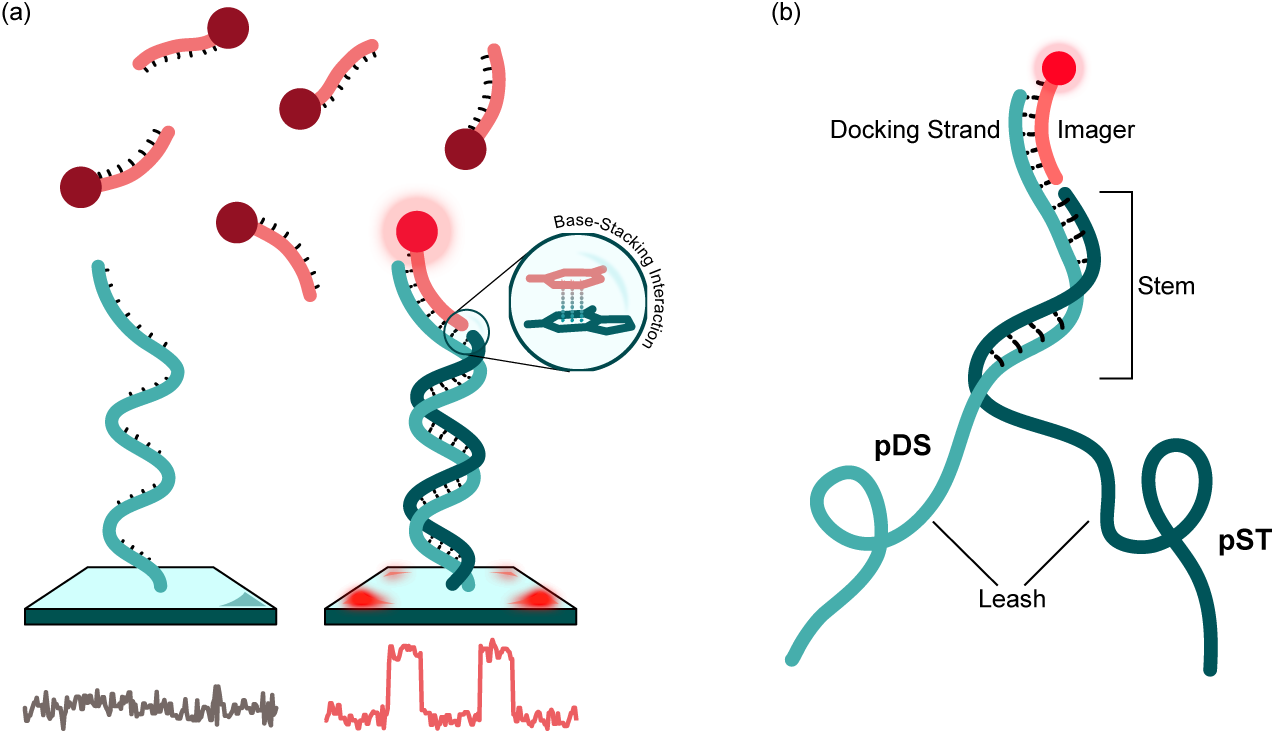
Design of Stack-pPAINT probe. (a) Base-stacking interactions facilitate detectable imager binding. (b) The split probe design includes pDS (proximity Docking Strand) and pST (proximity Stem).

In short, the docking strand in the Stack-pPAINT is constituted by two interacting DNA strands. First strand acts as regular docking strand of 6 nt in length but will not be sufficient on itself to allow detectable imager binding. The second strand upon hybridizing on docking strand creates the stem providing additional base-stacking interactions for the imager. Importantly, only when both strands are proximal to each other, we will detect the DNA-PAINT signal. Each strand can be used for encoding an individual biomolecule, enabling us to image the proximal molecules at high-resolution.

Each of the DNA strands are split into sections. The DNA sequence carrying the docking strand (pDS, proximity Docking Strand) is composed of a leash sequence, a stem sequence, and followed by a docking strand (Figure 1b). The leash sequence serves as a freely extendable spacer allowing for a greater area of exploration. The stem region works as the AND operator, hybridizing to the complimentary strand carried by the interacting partner (pST, proximity Stem) when present at proximity to one another (Figure 1b). This forms a platform for the imager strand to bind with both base-pairing and base-stacking interactions. The additional stacking interactions that arise from the recessed end of the stem are stabilizing in nature.^5^ This stabilization is tuned such that the presence of this interaction is essential for detectable binding events. In the absence of this interaction, the binding events would be extremely short lived thus would never be detected at the time scales of the camera’s frame rate and the extremely low signal-to-noise ratio that it would accompany.

### Molecular simulations highlight the interactions that drive proximity detection

To better understand the equilibrium conditions of Stack-pPAINT probe and study their relative stability across different designs, we carried out MD simulations using AMBER24^6, 7^. We created four unique all-atom models (System 1: Proximity probe, System 2: Proximity probe without fluorophore, System 3: Proximity Probe with continuous strand and no nick between imager and stem, System 4: Proximity probe without stem for stacking interaction; Supplementary Figure 1) to study the effect of nick, site at which stacking interactions occur, and fluorophore on the imager on microscopic configurations of proximity detection system (Figure 2a). To mimic the effect of distance constrains of the probe pair, we harmonically restrained O3’ and O5’ atoms of terminal nucleotide throughout the simulations with a spring constant of 1 kcal/mol/Å^2^. Two replicates of each all-atom models of DNA-dye constructs were simulated using NPT ensemble for 1 µs.

**Figure 2:**
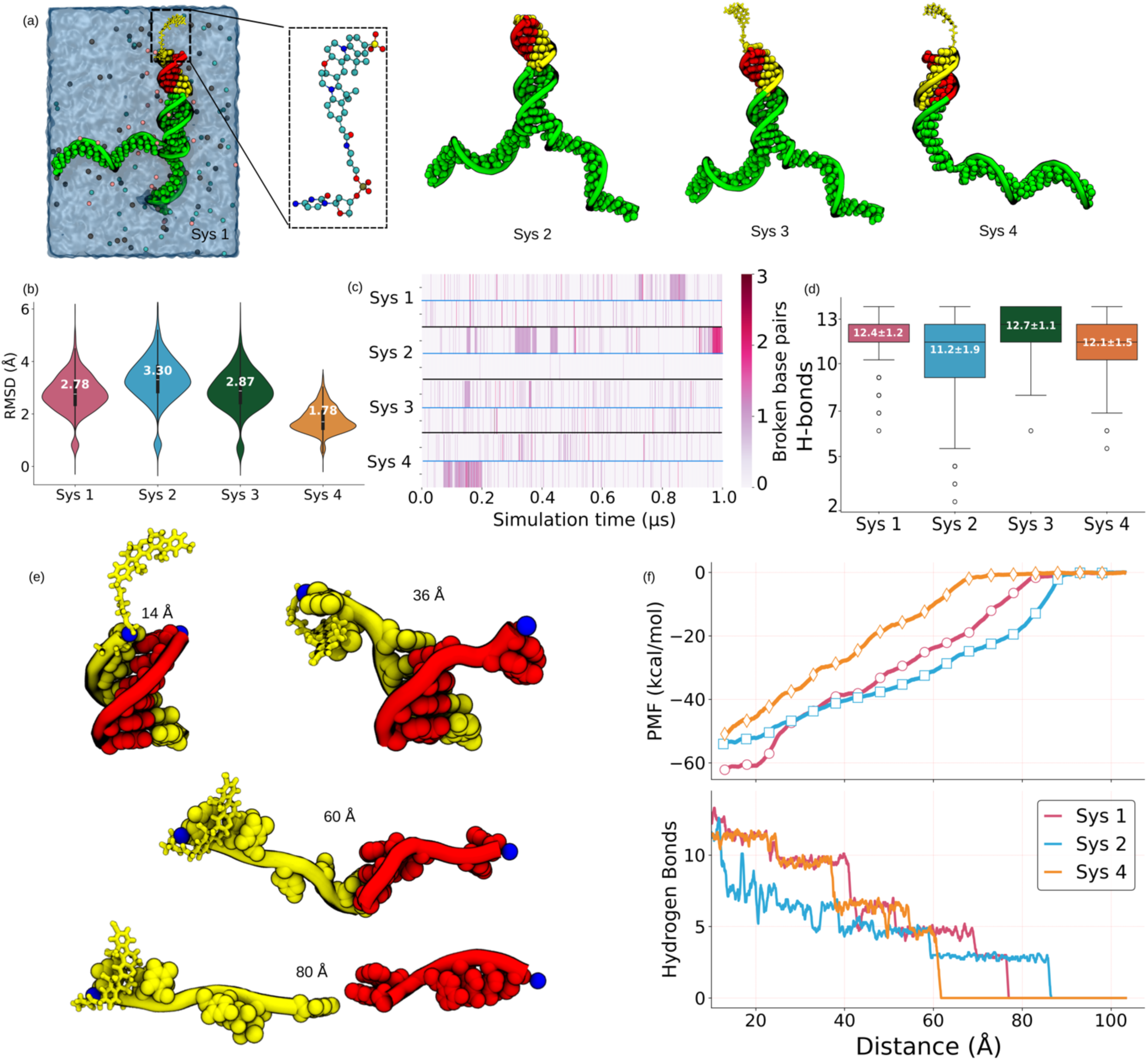
Equilibrium and advanced sampling MD simulations show the effect of different components on probe stability: (a) Left, a typical snapshot of the initial configuration (system 1) of dye conjugated DNA in solution. DNA atoms are shown using yellow (imager), red (hybridized with imager) and green (rest of the docking DNA) spheres. The volume occupied by water and ions is shown using transparent blue background. The dye molecule is shown is liquorish yellow. The covalent conjugation of the dye to DNA nucleotide (non-hydrogen atoms) is highlighted inside the dotted box. The atomic configuration of system 2 to 4 are shown without water for clarity. (b) RMSD distribution of dsDNA atoms (14 paired bases) with respect to initially built models. (c) Broken base pairs in the dsDNA region (14 bp long) as a function of simulation time for both the runs for various systems. (d) Boxplot showing the distribution of the total number of Watson-Crick hydrogen bonds between dsDNA. The simulation data from both the runs of respective system is concatenated for RMSD and hydrogen bonds shown in (b) and (d). (e) Instantaneous snapshot of the DNA illustrating various stages of ASMD. (f) (Top) Potential of mean force (PMF) and (bottom) number of hydrogen bonds between imager (yellow) and docking (green) strand as a function of the distance between oxygen atoms being pulled in ASMD simulations. The blue sphere highlights the terminal oxygens (O3’ of imaging strand and O5’ of docking strand) those were pulled apart.

Starting with the fully hybridized conformations (Figure 2a), the double stranded (dsDNA) region containing 14 paired bases (System 1, 2, and 3) or 6 paired bases (System 4) largely remains stable during microseconds long equilibrium simulation. The root mean square deviations (RMSD) of the dsDNA with respect to the initially built structure saturates within 3 Å (Figure 2b). Occasional breaking of two to three base pairs out of 14 was observed during the simulation (Figure 2c and 2d). As expected, the single stranded (14T) part of the models displays high flexibility adding significantly to RMSD of the entire DNA, 12Å-14Å (Supplementary Figure MDS3). We notice that the base pairs of the dsDNA regions remain stacked throughout the simulation (Supplementary Movie 1), the other geometrical parameters, like helical twist, rise etc. also remain close to the canonical B-DNA conformations (Supplementary Figure 2).

The analysis of equilibrium simulation trajectory shows that the nick in the backbone between imager and stem (System 1) does not add any structural deviations compared to the non-nicked structure (System 3). The terminal bases remain stacked for throughout simulation (Supplementary Figure 2), suggesting that stacking interactions may be enough for maintaining the stability of the DNA duplex despite one nick on the backbone. The RMSD (2.8 Å) and average number of broken base pairs (0.59) are nearly identical for nicked and the non-nicked structures (Figure 2b, 2c and 2d) affirming the strong base stacking interactions^8^.

The dye-DNA interactions are interesting^9^, yet relatively less explored aspects of the hybridization kinetics of nucleotides. In all the systems containing the dye molecule, Cy3B is covalently conjugated to the overhanging 3′ terminal nucleotide (DC3) (Figure 2a, Supplementary Figure 1) of imager and placed in an unstacked conformation at the beginning of simulation runs. During the simulations, the dye molecules exhibit a strong tendency to minimize solvent exposure by engaging in stacking interactions with the neighboring DNA bases likely driven by its polycyclic aromatic structure. Attributing to the strong stacking interaction of the dye to DNA, the dye containing systems show more structural stability compared to the non-dye DNA model (System 2) (Figure 2b, 2c and 2d). Principal component analysis of the backbone atoms from equilibrium MD simulations shows top three modes of conformational fluctuation in the DNA systems (Supplementary Figure 5) which corresponds to 60 % of the total motion.

To further understand the relative stabilities of the hybrids, we employed ASMD simulations where we pulled the imager away from the docking strands at a constant velocity of 0.5 Å/ns and measured the force-extension characteristics (Figure 2e and Supplementary Figure 6). The work done by this external force is converted to free energy difference (ΔG) or potential of mean force (PMF) of the hybridized versus non-hybridized states using Jarzynski equality^10^ (Figure 2f). During this 180 ns non-equilibrium simulations, (Supplementary movie SM2), the number of hydrogen bonds between the imager and docking strand goes from 33 to 0, marking their complete detachment (Figure 2e and 2f). The estimated free energy (ΔG) to dissociate the imager from the docker ranges between 50 to 75 kcal/mol. The higher values of ΔG as compared to expected from nearest neighbor model^11^ or experimentally measured free energy for similar systems^12^, could be due to ultrafast pulling velocities in simulations or due to notorious sampling problem. Nonetheless we believe the values qualitatively capture the trend correctly. The system 4 displays least stability in terms of ΔG which can be attributed to the missing stacking interaction between stem and imager (there is no stem). Comparing ΔG of system 1 and system 2, we find that that addition of dye stabilizes the system corroborating with the equilibrium simulations where extra stability from the dye was observed.

### Stacking interactions facilitate proximity detection

As an experimental platform, we utilized DNA origami-based nanostructures^13^ to showcase the usability of Stack-pPAINT approach^2, 3^ (Supplementary Figure 7). These modular structures allow for positioning of DNA strands in a myriad of ways at defined distances. We first designed staples where both the interacting partners emerge at distance of five nanometers from each other arranged in a 4×3 manner, where every pair is separated from one another by 20 nm (Figure 3a and 3b). Under conditions where both pDS and pST are present, clear 4×3 grids are visible using the imager for Stack-pPAINT docking strands. In the absence of the pST, there are no signals visible from the Stack-pPAINT docking strand thus indicating the efficiency of our probes in enabling signal detection only under proximity of the target pair (Figure 3c, 3d, and Supplementary Figure 8).

**Figure 3:**
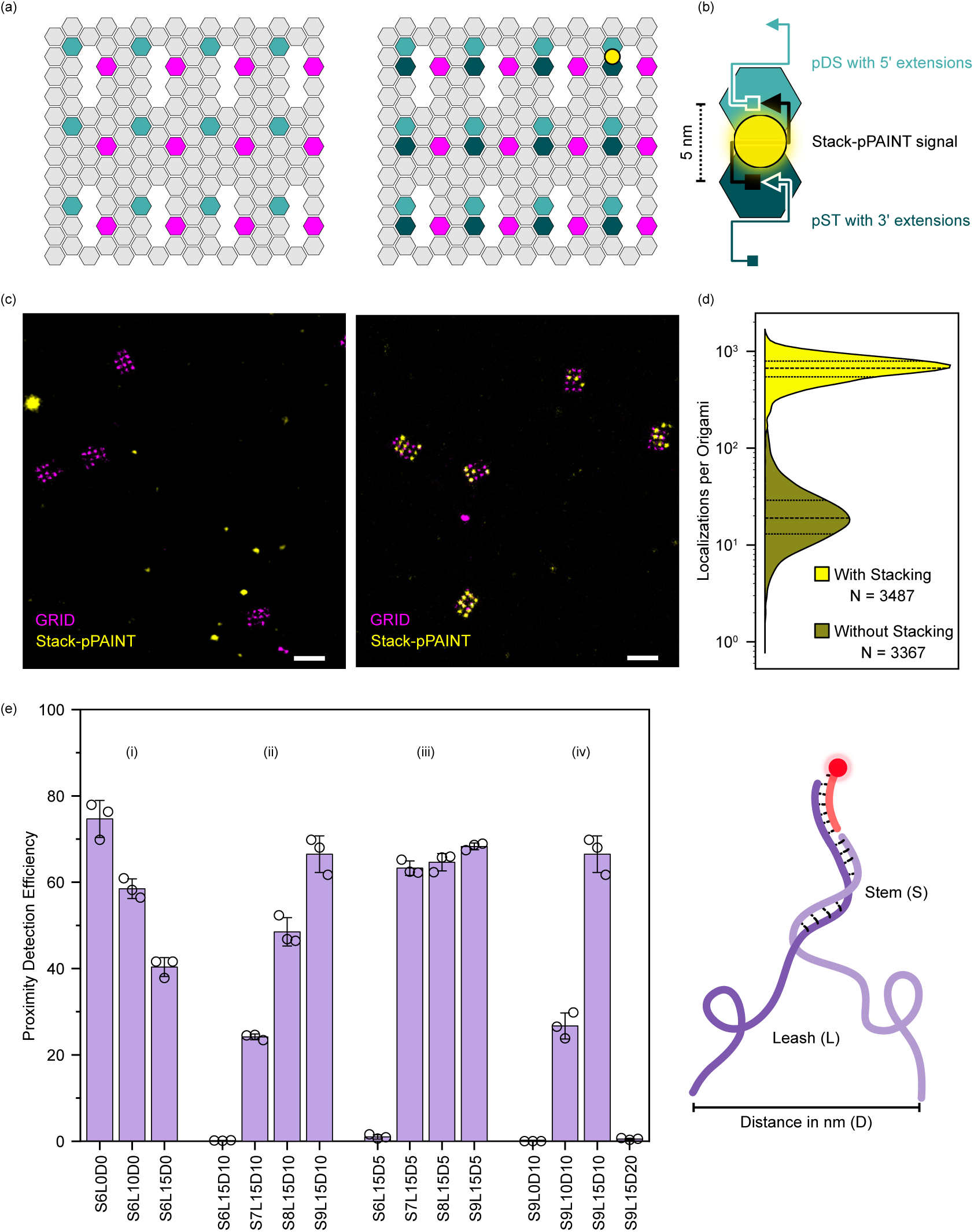
Stack-pPAINT efficiency: (a) DNA origami design for assaying Stack-pPAINT modality. pDS is present in both the conditions. pST is either absent (left) or present (right) in either experiment, highlighting the need for stacking interactions in stack-pPAINT. Direct extension docking strands allow for visualization of the origamis with an orthogonal imager (Magenta) (b) Staple design for stack-pPAINT and expected location of signal. (c) Signal of Stack-pPAINT (yellow) in the absence (Left) and presence (Right) of pST at a distance of 5 nm to pDS. Origami locations were detected using direct docking strand extensions (Magenta), (d) Distribution of the number of localizations per origami with and without stacking interaction (pST present or absent). (e) (i) Efficiency with increasing leash length (L) at 0 nm distance (D) with Stem (S) length of 6 nt; (ii) Efficiency with increasing stem length (S) at 10 nm distance (D) with 15 nt Leash (L); (iii) Efficiency with increasing stem length (S) at 5 nm distance (D) with a Leash (L) of 15 nt; (iv) Efficiency with increasing leash length at 10 nm distance (D) and non-specific interaction control at 20 nm distance (D). Schematic of Stack-pPAINT (right).

### Leash and stem lengths dictate the efficiency of proximity detection

To establish the optimal leash and stem lengths for efficient proximity detection at varying distances between the two targets, we used DNA origami structures as molecular breadboards. We place the pDS and pST at varying distances with varying stem and leash lengths to quantitively establish their detection efficiency (Supplementary Figure 9). We observe that at a closest distance where pDS and pST are next to each other (0 nm), we achieve the highest efficiency in the absence of any leash nucleotides. By increasing the leash nucleotides at this length scale, we see a drop in the efficiency, likely caused by steric hindrance imposed by the additional nucleotides present in the local vicinity (Figure 3e(i)). When the targets are placed 10 nm apart, an interacting partner pair with a six-nucleotide stem is insufficient to allow any detection of proximity. Stem lengths of seven or longer start showing clear proximity detection signals with a nine-nucleotides stem showing the highest efficiency. This can be attributed to the tug of war between the interaction strength of the base-pairing in the stem and the inherent tendency of ssDNA to collapse due to shorter persistence lengths (Figure 3e(ii)). A six-nucleotide stem would be entropically inefficient for such a weak duplex formation and would survive for shorter durations of time thus reducing the probability of stem formation and imager binding. A nine-nucleotide stem on the other hand is more stable which can overcome the entropic cost and thus would likely survive long enough to facilitate imager binding with stacking interactions. A similar but less pronounced effect is seen when the targets are placed 5 nm apart where a change from six-nucleotide to seven-nucleotide is sufficient to achieve almost maximum efficiency (Figure 3e(iii)). Upon testing how nine-nucleotide stems affect detection with different leashes where the targets are placed 10 nm apart, we observe that a 15-nucleotide leash is necessary and sufficient to obtain maximum achievable detection efficiency (Figure 3e(iv)).

To check if this combination of nine-nucleotide leash and 15-nucleotide stem facilitates non-specific target detection by facilitating non-proximity-based interactions, we placed the two targets 20 nm apart. Any signal from this condition can only potentially arise from the stable interactions between the two target pairs in solution prior to origami folding. We observe no detectable signal from this layout, prompting us to conclude that there aren’t any stable non-specific interactions occurring in solution that is driving the higher rates of proximity-detection (Figure 3e(iv)).

### Super-resolution imaging in cells using Stack-pPAINT

To extend this approach of proximity detection for use in cellular proximity detection, we labeled microtubule α-tubulin and β-tubulin subunits. For this, we used two different species of primary antibodies for labeling the individual subunits. Secondary antibodies of corresponding species were conjugated with pDS and pST. This brings a new hurdle where cooperative multi-strand interactions between the stem regions of the DNA could lead to non-proximity driven interactions between the secondary antibodies. This could lead to potential false detection of proximity where a single target is present at very high density. To circumvent this, (i) we conjugated secondary antibody to achieve a stochiometric average of three DNA strands per secondary antibody, (ii) we reduced the stem length from the most efficient number of nine nucleotides to eight nucleotides, where a majority of signal was still being detected. Next, we employed a rigorous chemical wash (30% DMSO in 1×PBS) to denature DNA hybridization to remove non-specifically bound antibodies via multivalent DNA hybridization. This step ensures to retain secondary antibodies directly bound to the primary antibodies but wash off secondary antibodies that were bound via DNA hybridization only (Figure 4a).

**Figure 4:**
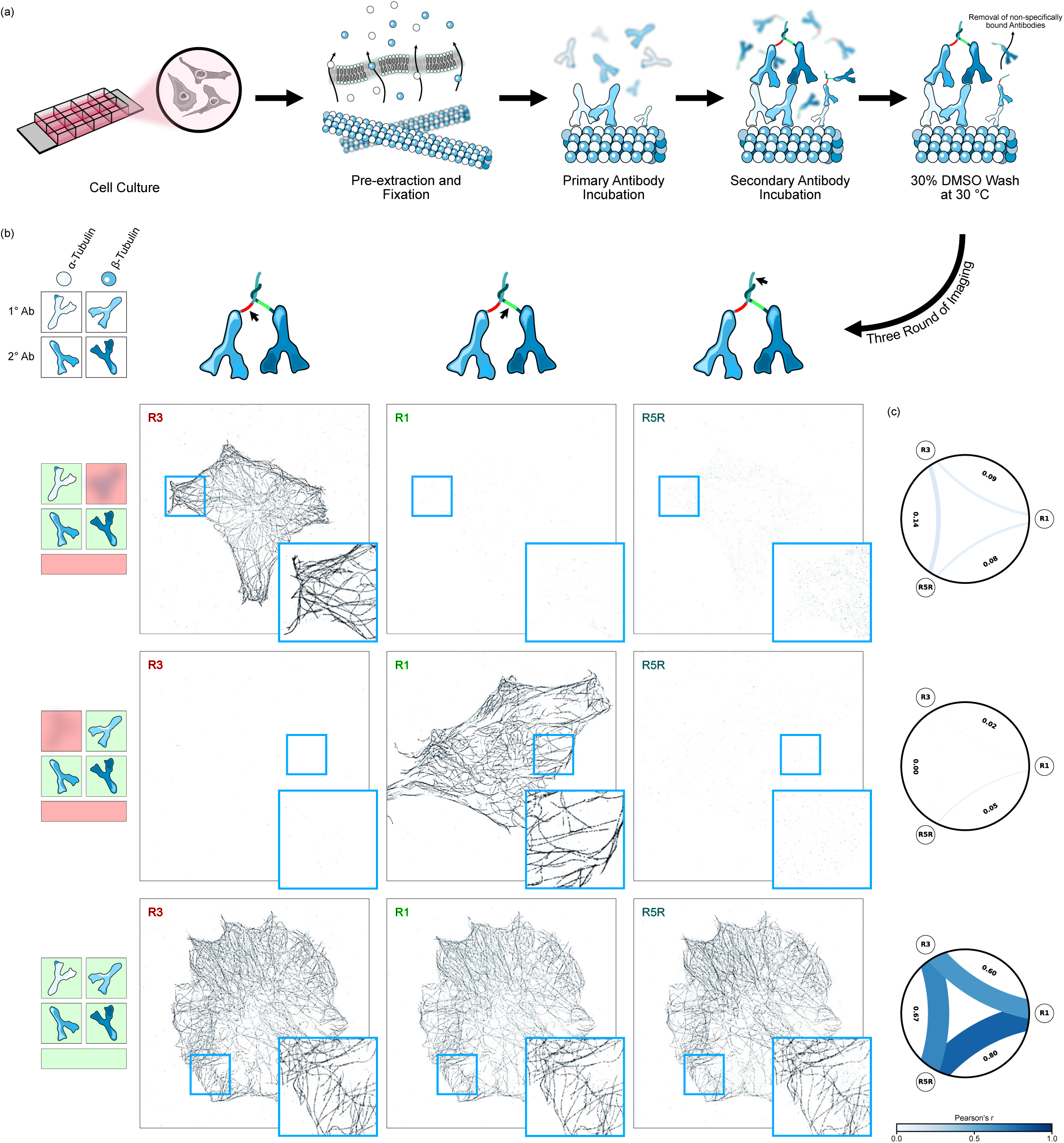
Stack-pPAINT in the cellular setup: (a) Workffow for microtubule staining by Stack-pPAINT antibodies. (b) Secondary antibodies have internal strands to visualize each secondary antibody species. Only in the presence of both the primary antibodies does the Stack-pPAINT R5R imager show signal. (c) Pearson’s r of the localizations between each imager across all three experimental conditions.

We were ourselves warned about the multivalent interactions of the 8-nt stem formed between two secondary antibodies, which could stabilize them even in the absence of either of the primary antibodies, potentially leading to non-specific signals. As a control, we first stained cells with either of the primary antibodies to ensure only one of the subunits are stained (Figure 4a). These served as negative controls where only one species of secondary antibody would have its target during staining. To directly assay the presence of the secondary antibody, we introduced orthogonal docking strands within the leash for pDS and pST. Under our stringent conditions – including washes with DMSO and fewer docking strands per antibody – we observe that, in cases where only one primary antibody was used, no signal from the Stack-pPAINT imager was detected, while signal from the secondary antibody in its cognate channel remained present (Figure 4b).

In contrast, when both primary antibodies were present, we observe clear signals from the Stack-pPAINT probe. The measured width of the microtubular network was approximately 60 nm, comparable to values reported by other super-resolution imaging approaches (Supplementary Figure 10). This demonstrates the capability of our probe to resolve cellular heterodimeric populations with high-resolution (Figure 4b). To further quantify the specificity of Stack-pPAINT, we assessed signal colocalization across the datasets where either primary antibody was omitted. Pearson’s *r* for the signal colocalization was very weak (below 0.1) in these controls, but strong (0.6-0.8) in the Stack-pPAINT condition (Figure 4c), clearly indicating the specific and proximity detection enabled by our approach.

## Conclusion

We designed a novel DNA-PAINT probe that can detect cellular proximity with high-resolution and specificity with the help of DNA base-stacking interactions. We fine-tuned DNA probes design and stacking energies required for detectable binding of the imager with the docking strand only when two strands are proximal. We performed all-atom equilibrium and pulling MD simulations to quantify the binding energetics, and the interactions between the pDS and pST parts of the probe and the imager. The molecular level simulations reveal that the additional stacking interaction in the presence of the fluorophore provide a considerable stabilization, thus allowing for longer and detectable binding. We further showcase the usability of Stack-pPAINT in the cellular setting by imaging the microtubular network made up of α-tubulin and β-tubulin monomers. Our probe design can be readily adapted to visualize heterodimeric interactions within the cell at high-resolution. Utilizing qPAINT^14^ along with Stack-pPAINT can further enhance the capabilities of this technique to extract quantitative information of heterodimeric interactions within cells.

## Methods

### General Simulation Methodology

All-atom MD simulations were performed using the MD simulation program AMBER24^6^, periodic boundary conditions and particle mesh Ewald (PME)^15^ method to calculate the long range electrostatics. Cuda enabled pmemd.cuda^16^ simulations were performed on RTX4080 machines. The Monte Carlo barostat^17^ with isotropic position scaling and Langevin thermostat^18^ with 1 ps^-1^ collision frequency were used to maintain the constant pressure and temperature in the system. AMBER force fields^19^ were used for describing the bonded and non-bonded interactions among DNA, water and ions. 8 Å cutoff was used to calculate van der Waals and short-range electrostatics forces. All simulations were performed using a 2 fs (femto-second) time step to integrate the equation of motion. SETTLE^20^ algorithm was applied to keep water molecules rigid whereas RATTLE^21^ algorithm constrained all other covalent bonds involving hydrogen atoms. The coordinates of the system were saved at an interval of 20 ps. The analysis and post processing the simulation trajectories were performed using VMD^22^ and CPPTRAJ^23^.

### All-atom model of Cy3B and its conjugation DNA backbone

Structure of Cy3B was taken from PubChem (CID: 44140555) and modified by adding amino linker – C3. The geometry of the molecule was subsequently optimized using Gaussian16^24^ followed by vibrational frequency calculations to verify the optimized structure as a true minimum by checking for imaginary frequencies using Density Functional Theory^25^. B3LYP/6-31+G level of theory was chosen to balance computational efficiency and to include polarization functions to improve electron distribution description. The keywords iop(6/33=2,6/42=6) directives enable finer control over natural bond orbital and ensure correct charge distribution for subsequent computer simulations. Antechamber^26^ program of AmberTools23^7^ was used to make the covalent link between dye and DNA, and subsequently prepare input files for LEaP from gaussian output by using RESP charge fitting. Figure MD1A describe the covalent link of Cy3B to DNA backbone.

### Building of DNA Nanostructure

The nucleic acid builder (NAB)^27^ module of AmberTools^7^ was used to generate models of DNA with exact same sequence used in experiments. Poly-thymine within DNA were introduced using mutagenesis plugin of PyMOL^28^ and the poly-thymine domain (31 bases on both strands) was further stretched and rotated in xLEaP to closely mimic the experimental setup. Mg^2+^ counterions were added to neutralize the charge of DNA. Additionally, K^+^ and Cl^-^ ions were added to obtain 150 mM concentration of KCl using LEaP module that creates a coulombic grid of 1 Å around the solute and places the ions at lowest electrostatic potentials^29^. Finally, all the systems were solvated in a rectangular TIP3^30^ waterbox ensuring 10 Å buffer around DNA. The all-atom topology and parameters were generated using xLeaP module of AmberTools^31^. ff99SB^32^ DNA forcefield parameters with bsc1^33^ modification were used to describe the bonded and non-bonded interatomic potentials of nucleotides. GAFF^34^ module of AMBERTOOLs was used to parameterize the force field of Cy3B. Four such all-atom models were created namely System 1: Proximity probe, System 2: Proximity probe without fluorophore, System 3: Proximity Probe with continuous strand and no nick between imager and stem, System 4: Proximity probe without stem. Each system contained about 110K atoms and measured 10 x 10 x 10 nm ^3^ (Supplementary Table 1).

### Equilibrium MD simulations

Thus, assembled systems underwent 2000 steps energy minimization using a steepest-descent algorithm to remove any bad clashes in the system. Following the energy minimization, the systems were gradually heated up to 300 K in 1 ns MD simulation with 2 fs time step. While heating, all non-hydrogen atoms of DNA and Cy3B were harmonically restrained having 1 kcal/mol Å^-2^ spring constants. Subsequently, we equilibrated systems for first 40 ns at 1 atm pressure and 300 K temperature to relax all water molecules and hydrogen atoms having same restrained atoms in heating at 1 kcal/mol and 0.1 kcal/mol. We constrained the O3’ and O5’ atoms of the docking stand and stem respectively in system 1-4 for whole simulation to mimic the immobilization to DNA origami surface (only O5’ in system 4 as there is no stem in this system) with a spring constant of 1 kcal/mol Å^-2^.

All systems were simulated with explicit water and ions at normal pressure (1 bar) and temperature (300 K) i.e. NPT ensemble. Two replicas of each system were simulated each lasting minimum 1 µs. Figure MD1(a) shows a typical snapshot of the systems at the beginning of the simulation. Supplementary movie SM1 shows the all-atom MD trajectories of the simulated systems, water and ions are not shown.

### Adaptive Steered Molecular dynamics (ASMD)

After equilibrating the DNA systems for 20 ns with non-hydrogen atoms of DNA restrained, we performed adaptive steered MD simulation (ASMD)^35, 36^ to study the unbinding of imager in all systems except system 3. We performed ASMD simulation with constant velocity of 0.5 Å/ns and a force constant (k_spring_) of 7.2 kcal mol–1 Å–2 on terminal oxygen atoms (O3’ of imager and O5’ of docking strand). These two atoms are highlighted in blue in figure MD1(f). We performed naïve ASMD^37^ in 9 stages pulling terminal oxygens from 14 Å to 104 Å, in each stage we ran 4 replicas. With the help of Jarzynski’s inequality^10^ we determined the trajectory having work done closest to the Jarzynski average which was used for the next consecutive step. ASMD is useful in sampling large conformational space of hybridization along the direction of reaction coordinate while minimizing the computational cost. Supplementary movie SM2 shows the steered MD trajectories of the simulated systems, water and ions are not shown for clarity.

### Buffers

10× Folding Buffer: 500 mM Tris-Cl pH 8.0 (Tris-Base, Sigma #77861; HCl, Fisher Scientific #29507), 125 mM MgCl_2_ (SRL #) and 10 mM EDTA (SRL #35888).

Buffer A: 10 mM Tris-Cl pH 8.0 and 100 mM NaCl (SRL #41721).

Buffer A+: 10 mM Tris-Cl pH 8.0, 100 mM NaCl and 0.05% v/v Tween 20 (Sigma #P9416-100ML). Buffer B: 50 mM Tris-Cl, pH 8.0, 10 mM MgCl_2_, 1 mM EDTA.

Buffer B+: 50 mM Tris-Cl, pH 8.0, 10 mM MgCl_2_, 1 mM EDTA, 0.05% v/v Tween 20.

Buffer I+: 50 mM Tris-Cl, pH 8.0, 10 mM MgCl_2_, 0.2 mM EDTA, 0.05% v/v Tween 20.

PEM Buffer: 80 mM PIPES (SRL #49159), 5 mM EGTA (SRL #62858), 2 mM MgCl_2_, pH 6.8.

Cytoskeleton Extraction Buffer: 0.25% Triton X-100 (Sigma #T8787-250ML), 0.1% Glutaraldehyde (EMS #16020) in PEM, pre-heated to 37 °C.

Cytoskeleton Fixation Buffer: 0.25% Triton X-100, 0.5% Glutaraldehyde in PEM, pre-heated to 37 °C. Quenching Buffer: 0.1 mg/ml of NaBH_4_ (Sigma #213462-25G) in 1× PBS.

Blocking Buffer: 3% w/v BSA (Sigma #A4503-50G), 0.05 mg/ml Salmon Sperm DNA (Invitrogen #15632011) and 0.25% v/v Triton X-100 in 1× PBS.

Antibody Incubation Buffer: 3% w/v BSA, 0.05 mg/ml Salmon Sperm DNA, 0.02% v/v Tween-20 and 1 mM EDTA in 1× PBS.

Buffer C: 1× PBS supplemented with 500 mM NaCl.

### Flow Cell Preparation

We prepared slides and coverslips for microscopic imaging of DNA origami nanostructures as described earlier.^5^ Five pairs of holes were drilled in slides (VWR #631-1550) using a diamond head drill bit (Meisinger #801-009-HP) on a drill gun (DigitalCraft #LRUXOR) mounted on a drill stand (Dremel #220-01 Workstation™). Slides and coverslips (Marienfeld #0107222) were washed with 5% v/v dish washing detergent and thoroughly rinsed with Milli-Q® water followed by 70% ethanol and once again with Milli-Q®. A double-sided tape (3M Scotch #136D MDEU) was sandwiched between slide and coverslip to assemble flow cells. Using quick setting epoxy (Araldite Klear), the open ends were sealed. All imaging experiments using DNA origami nanostructures were done using these flow cells.

We used tubing attached to the drills for the exchange of buffers and imagers in multiplexed origami imaging.

### PEGylation of slides for Efficiency Measurements

To minimize non-specific sticking of the imager strand on the glass surfaces, we utilized PEG passivated slides. PEGylation was performed as previously reported^5^. In brief, coverslips and drilled slides were washed with 5% dishwashing detergent prior to PEGylation. In case the slides are being reused, they are first sonicated (Bandelin # Sonorex Super RK 255 H) in Acetone (Supelco #1070212521) for 20 minutes. Slides and coverslips were then rinsed thoroughly with MilliQ® water then placed on a home built Teflon holder for further steps. The slides and coverslips were then sonicated in MilliQ® water for 5 minutes in a glass beaker where the Teflon holder is immersed. MilliQ® water was then replaced with 1 M KOH (SRL #84749) and sonicated for a minimum of 30 minutes. MilliQ® water was then used to thoroughly rinse the slides and coverslips for at least three times.

Piranha etching was then performed by adding about 225 ml of H_2_SO_4_ (Supelco #1934002521) to the empty glass beaker followed by 75 ml of H_2_O_2_ (Qualigens #Q15465). Caution is taken as this highly reactive mixture is dangerous and hazardous. The slides are etched in this solution for 30 minutes or until the reaction reaches temperatures below 50 °C.

Slides and coverslips are then rinsed with MilliQ® water thoroughly to remove all traces of piranha solution. Slides are then rinsed thrice with Methanol (Supelco Emparta #1070182521) and sonicated in methanol for about 20 minutes.

Next, aminosilanation is performed by adding the aminosilanation mixture containing 25 ml of APTES ((3-Aminopropyl)TriEthoxySilane) (TCI #A0439), 12.5 ml of Acetic Acid (Supelco #1930022521), and 262.5 ml of methanol prepared in a dedicated flask stored with methanol before and after every use; into the glass beaker containing the Teflon holder carrying slides and coverslips. Aminosilanation is performed for a maximum of 15 minutes following which the mixture is replaced with methanol at least thrice followed by MilliQ® water thrice. Slides and coverslips are then dried using compressed nitrogen gas.

These aminosilanated slides and coverslips are then passivated using mPEG-SVA (Layson Bio #mPEG-SVA-5000) and Biotin-PEG-SVA (Laysan Bio #Biotin-PEG-SVA-5000) dissolved in 0.5 M NaHCO_3_ (Sigma #S5761) in a 4:1 ratio. 120 mg of mPEG-SVA and 30 mg of Biotin-PEG-SVA is dissolved in 960 µl of 0.5M NaHCO_3_. 60 µl of this solution is added to each slide and a dried coverslip is laid on it, sandwiching the added mixture. This setup is left overnight in a humidified box stored in a dark chamber.

The slides and coverslips are then rinsed thoroughly with MilliQ® water and then dried with compressed nitrogen gas before storing each pair in a 50 ml centrifuge tube ensuring that the PEG passivated slides are facing away from each other. The centrifuge tubes are then vacated and filled with inert nitrogen gas, sealed, and stored at -20 °C for up to 1 month.

Prior to use, the centrifuge tubes are placed and room temperature and are opened only after the temperature equilibrates. Flow cells are then prepared on these slides as mentioned above.

### DNA Origami Nanostructure Folding

All the staple sequences (Supplementary Table 2, 3 and 4) were generated using Picasso Design^38^ based on required design of DNA origami structures. All the DNA origami nanostructures were folded using an M13mp18 ssDNA (Bayou Biolabs #P107) scaffold. Origami structures were anchored to the surface via biotin-labeled oligonucleotide staples at 5′-end (Supplementary Table 2). Typically, we folded a 10 nM of scaffold DNA in a 30 µl of 1× folding buffer folding reactions with 100 nM of biotin and blank staples, 1 µM of staples extended with docking strands. The mix was incubated at 80 °C for five minutes before a stepwise cooling of 0.1 °C per every five seconds till room temperature in a thermocycler (Applied Biosystems™ ProFlex™ PCR System #4484073).

Excess staples from the folded origami nanostructures were removed using concentrator centrifugal spin filters (Sartorius Vivaspin® 500 #VS0132). The filters were wetted with Milli-Q® water and spun at 3000 ×g to remove water which was followed by 500 µl of 1× folding buffer. Freshly folded origami mix was diluted to 450 µl and applied to the column in 1× folding buffer and spun at 3000 ×g until the volume reached to below 50 µl. Another 450 µl of 1× folding buffer was added to the resultant, and this filtration was repeated three times to get rid of the excess staples. The origami structures were then collected and stored at -20 °C until use.

Each blank staple and biotin staple (Supplementary Table 2) was added in 10 molar excess to the scaffold. Staples for grid with direct extensions, 20 nm grid stack-pPAINT, and efficiency measurements were added in 100 molar excesses to the scaffold. Each blank staple is replaced by the corresponding grid, stack-pPAINT or efficiency staple as mentioned in Supplementary Table 3 and 4.

### Imager Fluorophore Conjugation and Purification

Fluorophore conjugation of DNA strands for preparing imager strands were done as per the published protocol.^5^ The imager sequences (Merck) were modified with amine at their 3′-end (Supplementary Table 5) and dissolved in Milli-Q® water.

Cy3B-MonoNHS-Ester (Cytiva #PA63101) was dissolved in DMSO (Sigma #D8418) at 13 mM concentration and Atto647N-MonoNHS-Ester (Sigma #18373-1MG-F) was dissolved at 11.8 mM concentration. Aliquots were stored at -20 °C. A 15 nanomoles of DNA in15 µl of 1× PBS and 0.1 M NaHCO_3_ was mixed with five-fold excess of Cy3B-MonoNHS-Ester or Atto647N-MonoNHS-Ester. The reaction was run for overnight at 4 °C while gently shaking.

Excess fluorophore and the unconjugated DNA oligonucleotides were separated from the conjugated product using reverse phase HPLC (Agilent 1260 Infinity Series II, equipped with a quaternary pump system). An oligo column (Phenomenex #00B-4442-E0 Clarity® 5 µm Oligo-RP, 50 mm × 4.6 mm) was used for all the purification. The purification processes involved gradient flow of three solvents which included solvent A (0.1M TEAA + 5% ACN), solvent B (0.1M TEAA + 50% ACN) and solvent C (100% ACN). The fluorophore-conjugated DNA was then lyophilized, dissolved in Milli-Q®, and then stored at -20 °C for until its usage.

### Sample Preparation for DNA Origami Nanostructure Imaging

Microfluidic flow cells were assembled as described earlier.^38^ First, a 10 to 15 µl of 1 mg/ml Biotinylated Bovine Serum Albumin (Sigma #A8549) diluted in Buffer A was incubated inside the channels. Next, the channel was washed Buffer A+. A 10 to 15 µl of 0.2 mg/ml of Neutravidin (Sigma #31000) dissolved Buffer A was incubated for around a minute. Excess neutravidin was washed with Buffer A+ thoroughly. Prior to imaging, the channel was washed with Buffer I+. Around 250 pM origami nanostructures diluted in Buffer I+ were introduced inside the flow cell and incubated for around 20 minutes. Unbound DNA origami structures were removed by washing with 600 – 800 µl of Buffer I+.

Gold Nanoparticles (Sigma #753688-25ML) were introduced into flow cell at a dilution of 1:10 in buffer I+ for 10 minutes and washed with 600 – 800 µl of Buffer I+ before performing imaging.

### Cell Culture

MEFs (Murine Embryonic Fibroblasts) were received as a generous gift from Dr. Kesavardana Sannula’s lab. They were grown in Dulbecco’s Modified Eagle Medium (DMEM Gibco #11995081) supplemented with 10% Fetal Bovine Serum (FBS Gibco #10270106) and 100 U/ml of Penicillin and 100 µg/ml of Streptomycin (Gibco #15140122) at 37 °C with 5% CO_2_. Cells were periodically maintained and split when they reached ∼70% confluency. For DNA-PAINT imaging experiments, cells were trypsinized with 0.05% Trypsin-EDTA (Gibco #25300054) for 30 seconds to a minute at 37 °C. Cells were then resuspended in DMEM, centrifuged at 300 ×g at room temperature for 4 minutes and the supernatant was discarded. The pellet was then resuspended in 1 ml of DMEM and Countess™ 3 Automated Cell Counter (Thermofischer) was used to count the number of cells. 4000 cells were then seeded on an 8-well cover glass bottom chamber slide (CellVis #C8SB-1.5H) in DMEM. Cells were then grown overnight in the incubator.

### Antibody Conjugation and Staining

Antibody conjugation was performed as mentioned previously. In short, anti-mouse donkey (Jackson Immunoresearch #715-005-150) and anti-rabbit donkey (Jackson Immunoresearch #711-005-152) were used in all experiments (Supplementary Table 6). 100 µg of antibody was used for each conjugation reaction. Antibodies were buffer exchanged using 50 kDa spin filter (Sartorius VivaSpin® #VS0132) at 12000 ×g at 4 °C into 1× PBS at a concentration >1.5 mg/ml. 5 molar excess of DBCO-PEG4-NHS Ester crosslinker (Jena # CLK-A134-10) which was dissolved in DMSO at a concentration of 1 µg/µl was added to the antibodies and incubated under a mild shaking condition on a shaker (Eppendorf #5382000023) for 2 hours at room temperature. The crosslinker was then separated using the same spin filters using 1× PBS with five rounds of repetition to dilute out the crosslinker. A conjugation ratio of about 1:3 for Antibody:DBCO is obtained. 5 molar excesses of Azide conjugated DNA strands were added to the respective antibodies (Supplementary Table 7) and incubated at 4 °C under a mild shaking condition on a shaker. Excess DNA was diluted out using similar steps as mentioned above.

### Cell fixation and staining

Cytoskeleton Extraction Buffer and Cytoskeleton Fixation Buffer were freshly prepared in PEM buffer prior to fixation. The cells grown on the glass bottom chamber slides were first pre-extracted with pre-warmed (37 °C) cytoskeleton extraction buffer for 30 – 45 seconds followed by fixation with pre-warmed (37 °C) cytoskeleton fixation buffer for 10 minutes inside the incubator (37 °C). Freshly made quenching buffer was then used to quench any free Glutaraldehyde for 7 minutes. The samples were then washed thrice with 1× PBS and stored at 4 °C for further processing. Blocking and Permeabilization is performed simultaneously with blocking buffer for 45 minutes to 1 hour. Primary antibodies (Supplementary Table 6) were incubated with cells in antibody incubation buffer overnight at 4 °C following which wells were washed with 1× PBS thrice. Similarly, secondary antibodies were incubated for 45 minutes at room temperature and washed with 1× PBS thrice. Cells were then incubated with a solution containing 30% DMSO in 1× PBS for 30 minutes at 30 °C. This step ensures all non-specific interactions mediated by the DNA molecules on the secondary antibodies are abolished. Cells were then washed with 1× PBS thrice.

### Stack-pPAINT Imaging and reconstruction

Both DNA origami structures and cellular samples were imaged on a Nikon Ti2E microscope body mated to a motorized H-TIRF unit and equipped with a Perfect Focus System (PFS). An oil immersion high NA TIRF lens (Nikon #Apo SR TIRF 100×, 1.49 NA, oil immersion) was used for TIRF (Total Internal Reflection Fluorescence) imaging along with the L6Cc laser combiner from Oxxius Inc, France for 561 nm and 640 nm excitation laser and a Teledyne Photometrics PRIME BSI sCMOS camera set at 16-bit readout sensitivity and 2×2 – pixel binning.

A 20× PCD (Protocatechuate 3,4-dioxygenase; Sigma #P8279-25UN) was prepared in 50 mM NaCl, 1mM EDTA, and 100 mM Tris-Cl, pH 8.0, and 50% glycerol to a concentration of 6 µM. The stock PCD was then divided into 10 µl aliquots and stored at -20 °C for future use.

A 40× PCA (Protocatechuic Acid / 3,4-Dihydroxybenzoic acid; Sigma #37580-100G-F) was made by dissolving 154 mg of PCA in 8 ml of Milli-Q® and 3 M NaOH (SRL #96311) was added dropwise and stirred until PCD completely dissolved. Volume was made up to 10 ml and 10 µl aliquots were stored at -20 °C for future use.

A 100× Trolox (6-hydroxy-2,5,7,8-tetramethylchroman-2-carboxylic acid; Sigma #238813-1G) was prepared by dissolving 100 mg of Trolox in 430 µl of methanol, 345 µl of 1 M NaOH and 3.2 ml of Milli-Q®. The solution was stored as 20 µl aliquots at -20 °C for future use.

For DNA origami structure imaging, 100 µl of imaging buffer composed of 1× PCD, 1× PCA and 1× Trolox was prepared in buffer I+. Imaging was performed with the 640 nm laser prior to 561 nm laser, both under TIRF illumination.

For cellular imaging, 200 µl of imaging buffer composed of 1× PCD, 1× PCA and 1× Trolox was prepared in buffer C. Imaging was performed with the 561 nm laser for all three rounds of imaging with an Exchange-PAINT^39^ modality. Between each round, sufficient wash with 1× PBS was until no localizations were observed in the absence of the imager.

Imager concentrations and imaging parameters are outlined in Supplementary Table 8.

Localizations were reconstructed and viewed using the Picasso^38^ (https://github.com/jungmannlab/picasso) suite of software. Cellular localizations were filtered to reduce noise based on photons, sx, and sy of the localizations. Filtering parameters are mentioned is Supplementary Table 9.

### Stack-pPAINT efficiency measurement

DNA origamis nanostructures were picked based on the signal from the direct extension grids with a pick size of 1 pixel. Signal from the Stack-pPAINT channel was extracted from each picked location. The percentage of picks showing signal (minimum of 5 localizations within the pick area) from the Stack-pPAINT channel was calculated and plotted.

### Pearson’s r measurement for Stack-pPAINT

All three channels of signal from cellular localizations were aligned using RCC on Picasso Render. Localizations were binned into pixels of 65 × 65 nm. Number of localizations for each pixel was tabulated, and Pearson’s *r* was calculated across each set and plotted.

## Author Contributions

A.B. performed experiments, analyzed data, and wrote the manuscript; S. Y. perform MD simulations, analyzed data, and wrote the manuscript. V.S.V performed experiments and analyzed data; S. K. performed experiments; S. A. provided lab resources and supervised S.K.; H. J. designed simulations study, analyzed data and supervised S. Y., and wrote manuscript; M.G. conceived and supervised the study, interpreted data, and wrote the manuscript. All authors reviewed and approved the manuscript.

## Acknowledgements

We acknowledge Department of Science and Technology, Ministry of Science and Technology, India DST-FIST Program funded Central Facility, Department of Biochemistry, IISc. This work has been supported by DBT/Wellcome India Alliance intermediate fellowship (IA/I/21/2/505928) to M.G. and the Department of Biotechnology (BT/PR40186/BTIS/137/3/2020) to M.G. We greatly acknowledge the support from Max-Planck Institute of Immunology and Epigenetics Freiburg, Germany in terms of partner group to Mahipal Ganji lab. A.B. acknowledges the support from the Prime Minister’s Research Fellowship (PMRF), Ministry of Education, Government of India. This work has also been supported by Anusandhan National Research Foundation fellowship (SRG/2022/002109) and the DST Inspire faculty fellowship (IFA20-PH-256) by the Govt of India to H.J. Supercomputing time was provided by PARAMSEVA through national supercomputing mission of Govt of India. S.Y. acknowledges the support received from Ministry of Education of Govt of India for her research fellowship. S.K. acknowledges the support received from Council for Scientific and Industrial Research, Ministry of Science and Technology, Government of India.

