## Supplementary Figures, Texts and Movies for "Proximity-based super-resolution imaging enabled by DNA base-stacking interactions"

#### Supplementary Figure 1

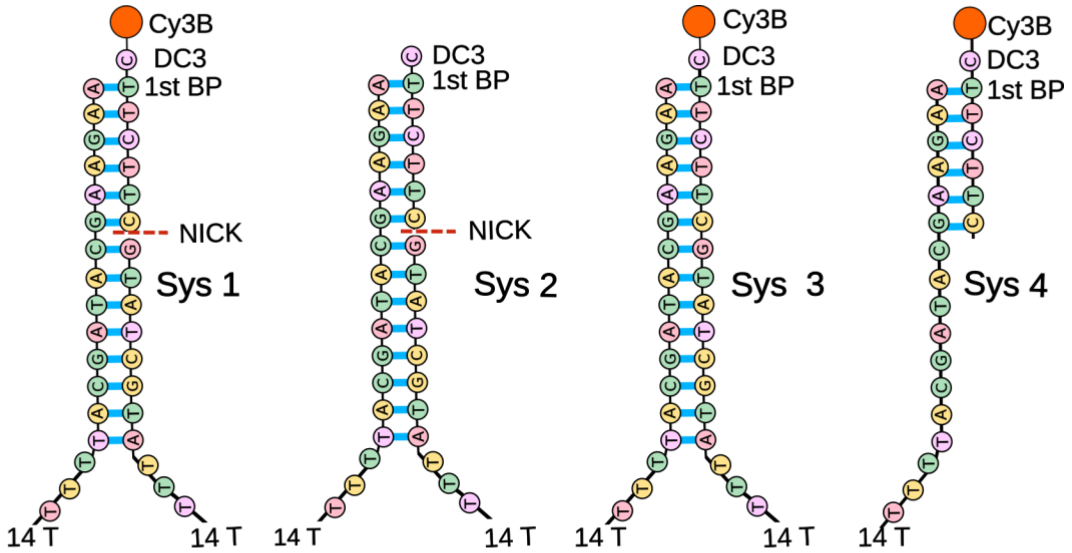


***Supplementary Figure 1: Schematic illustration of five all-atom models.*** *(i) System 1 comprises of a 7 base long imager bound to cy3B fluorophore along with both pDS and pST. (ii) System 2 lacks the fluorophore on the imager. (iii) System 3 consists of the entirety of system 1 but lacks the nick, making the imager and pDS a continuous strand. (iv) System 4 lacks the pST strand thus mimicking the condition where there lacks any proximity between pDS and pST. (v) System 5 has a 6 base long imager instead of a 7 base long imager.*

Supplementary Figure 2

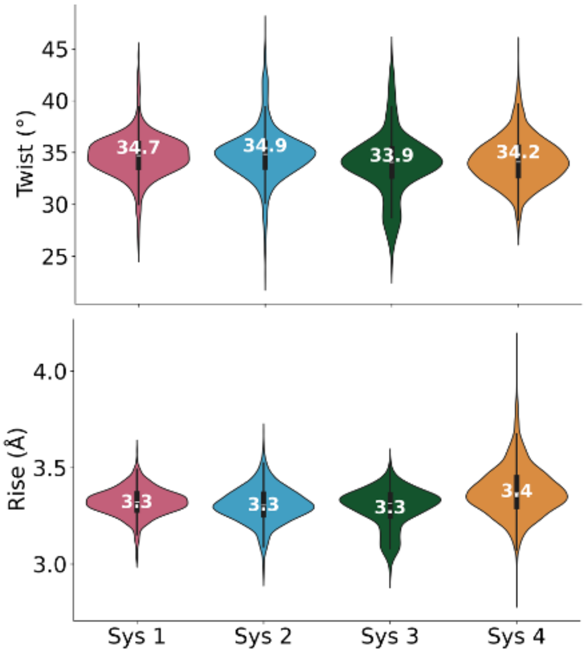


***Supplementary Figure 2: Helical parameters of DNA during simulations:*** *Probability distribution of average helical twist (top) and average helical rise (bottom) for the double stranded part of all atom models. Fourteen paired bases were taken in consideration for Sys 1,2,3,5 and six paired bases were considered for System 4. These geomatical parameters were averaged for entire simulations trajectory using 'nastruct' module of CPPTRAJ which utilizes the standard reference frame proposed by Olson et al*[1].

Supplementary Figure 3

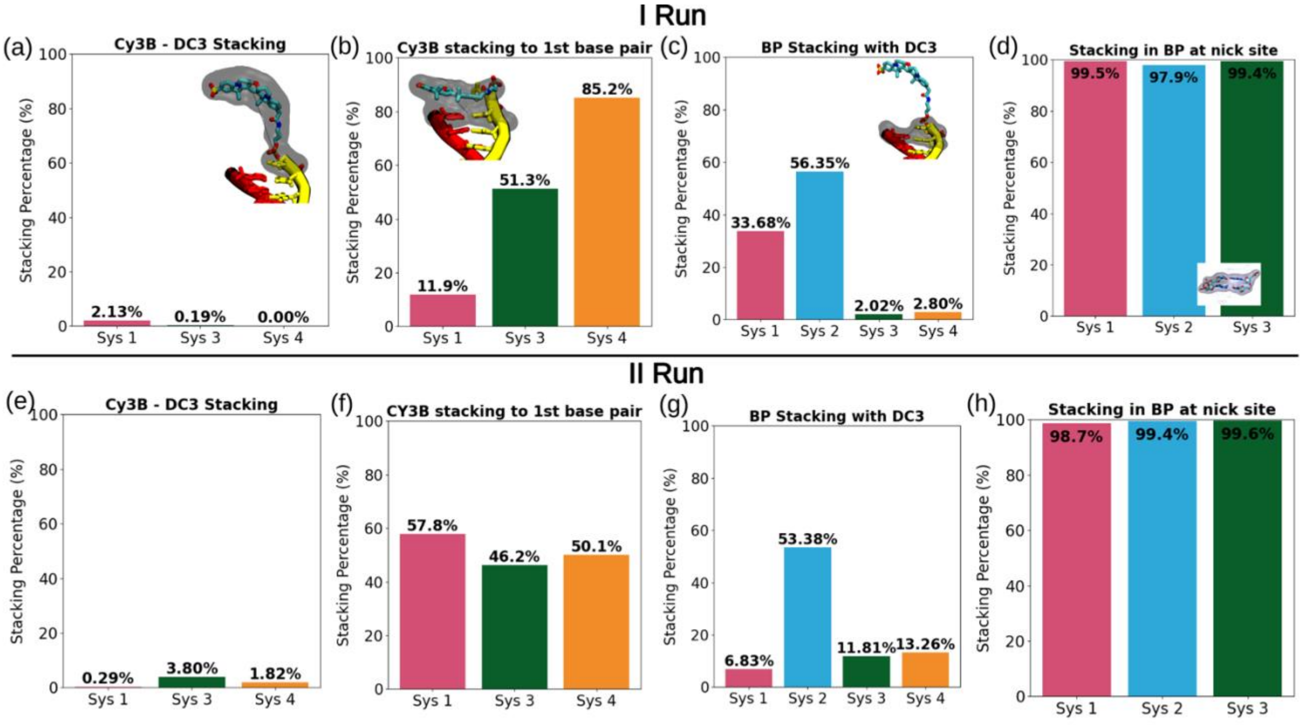


***Supplementary Figure 3: Stacking percentage within bases and fluorophores for two replica runs.*** *(a),(e) Cy3B-DC3: stacking of ring atoms of Cy3B with terminal base Cytosine for 3 systems containing dye and DC3. (b),(f) Cy3B with first base pair: Stacking of Cy3B with first base pair for four systems containing dye. (c),(g) BP stacking with DC3: stacking between terminal base, cytosine with immediate next base pair or first base pair. (d),(h) Stacking at nick site: Stacking interactions between base pairs immediately flanking the nick site. Highlighted region with black color shows representative residues under study.*

Supplementary Figure 4

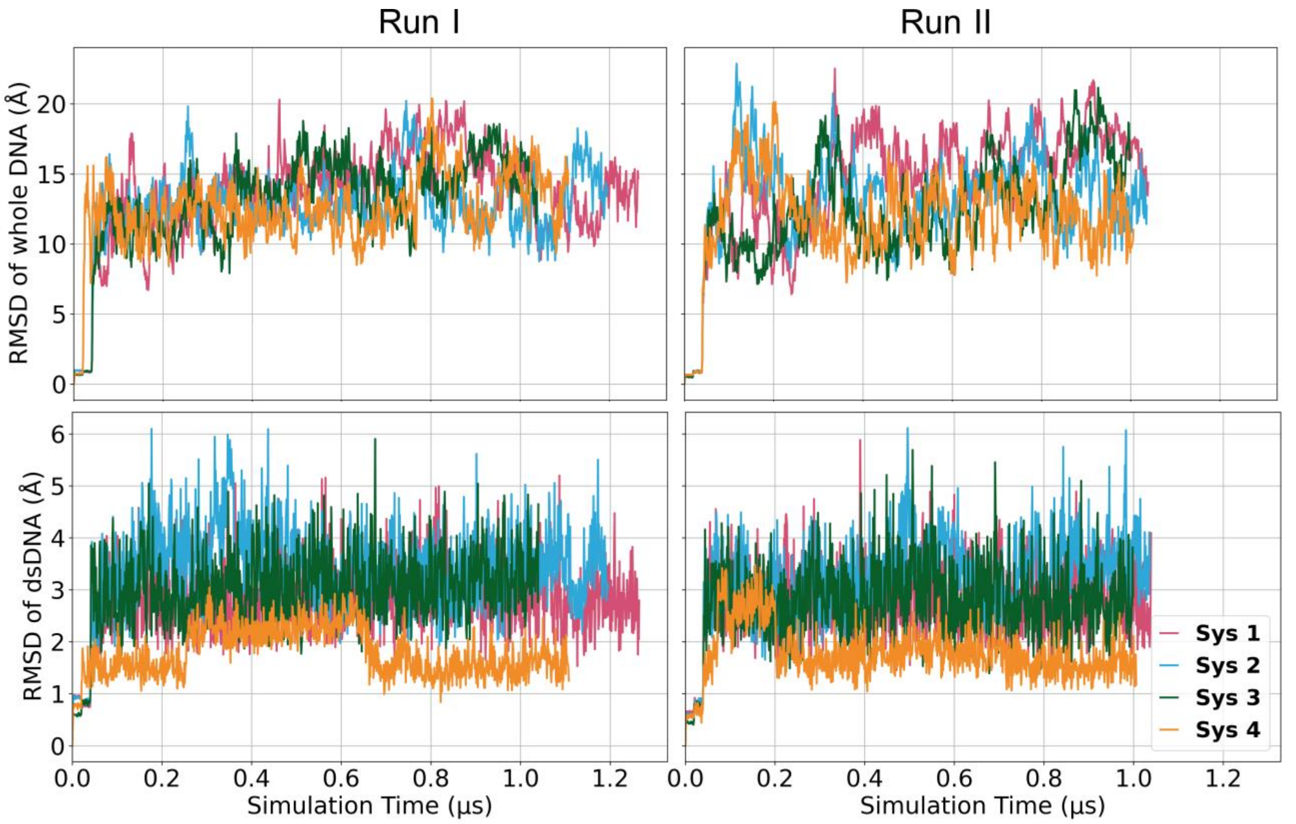


***Supplementary Figure 4:*** *RMSD of the whole DNA models (top row) and only dsDNA region (bottom row) in run 1 (left) and run 2 (right) with respect to the initially build structures as a function of simulation time.*

Supplementary Figure 5

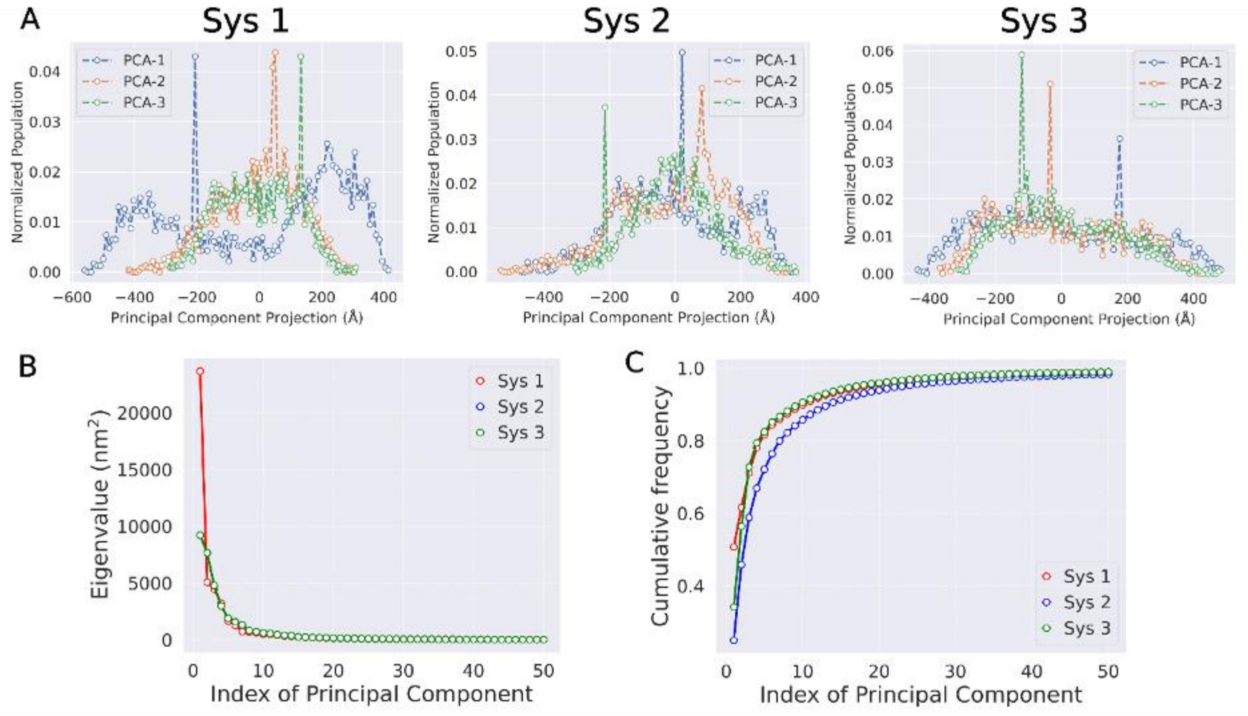


***Supplementary Figure 5: Principal Component Analysis (PCA) of DNA dynamics across three systems.*** *(a) Projection of trajectories along the first three principal components (PC1, PC2, and PC3) for each system, showing the normalized population distribution. (b) Eigenvalues corresponding to the first 50 principal components. System 1 shows a steep drop, indicating that the majority of the motion is captured by the first few components, while Systems 2 and 3 display more gradual decay. (c) Cumulative variance explained by successive principal components. System 1 reaches ~90% variance with fewer components compared to Systems 2 and 3, suggesting a more constrained motion dominated by fewer collective modes.*

Supplementary Figure 6

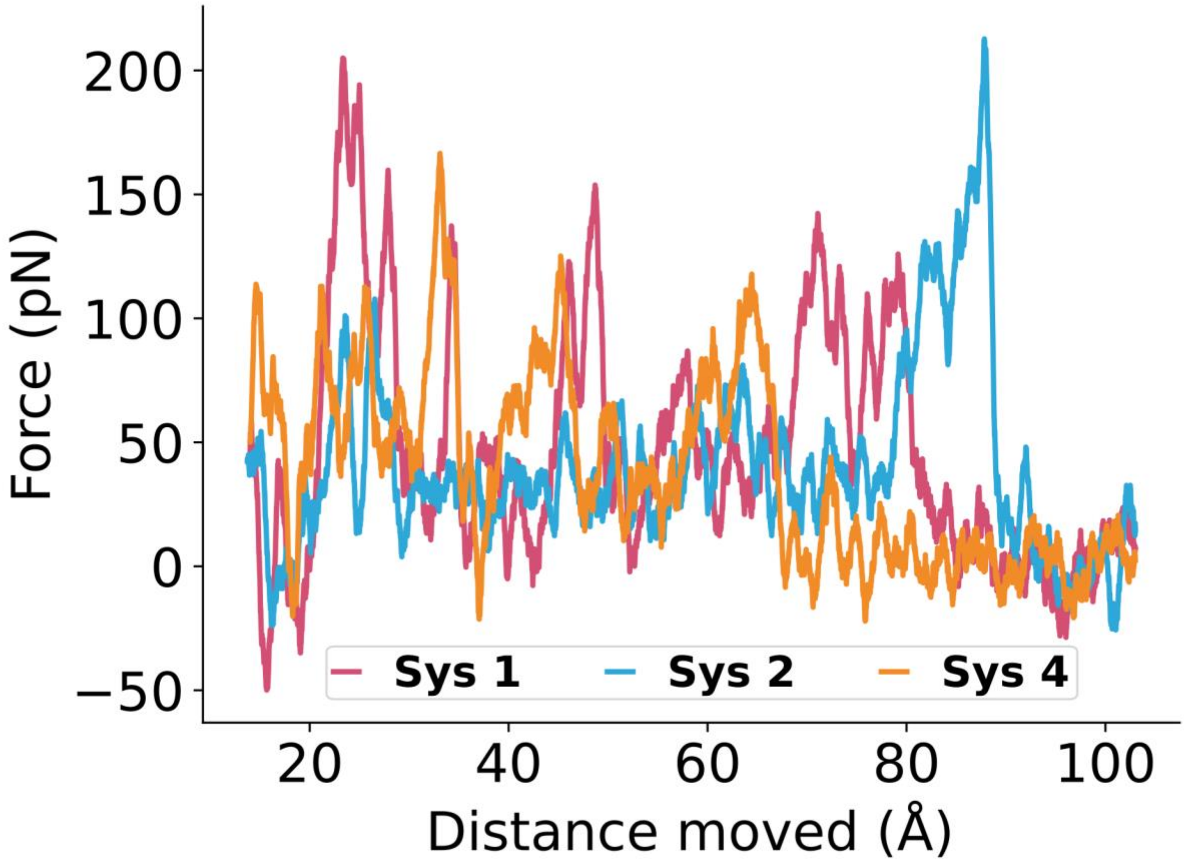


***Supplementary Figure 6: Force vs displacement graph for ASMD simulations.*** *Moving average of the raw force obtained during steered molecular dynamics (ASMD) simulations as a function of the distance moved (reaction coordinate) for systems.*

Supplementary Figure 7

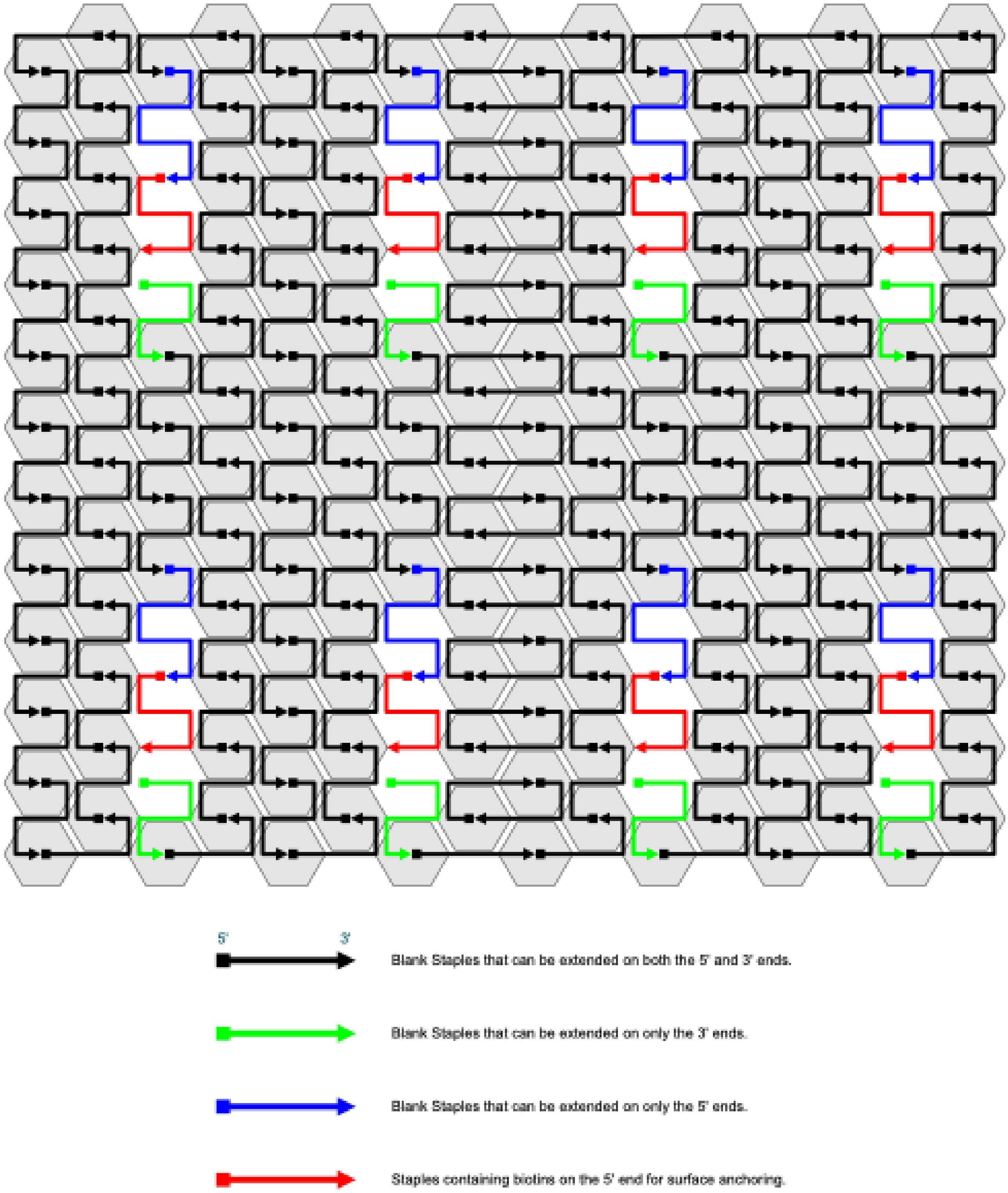


***Supplementary Figure 7: Staple outline:*** *Modified Rothemund's Rectangle[2] Origami outlining all the staples involved in folding.*

Supplementary Figure 8

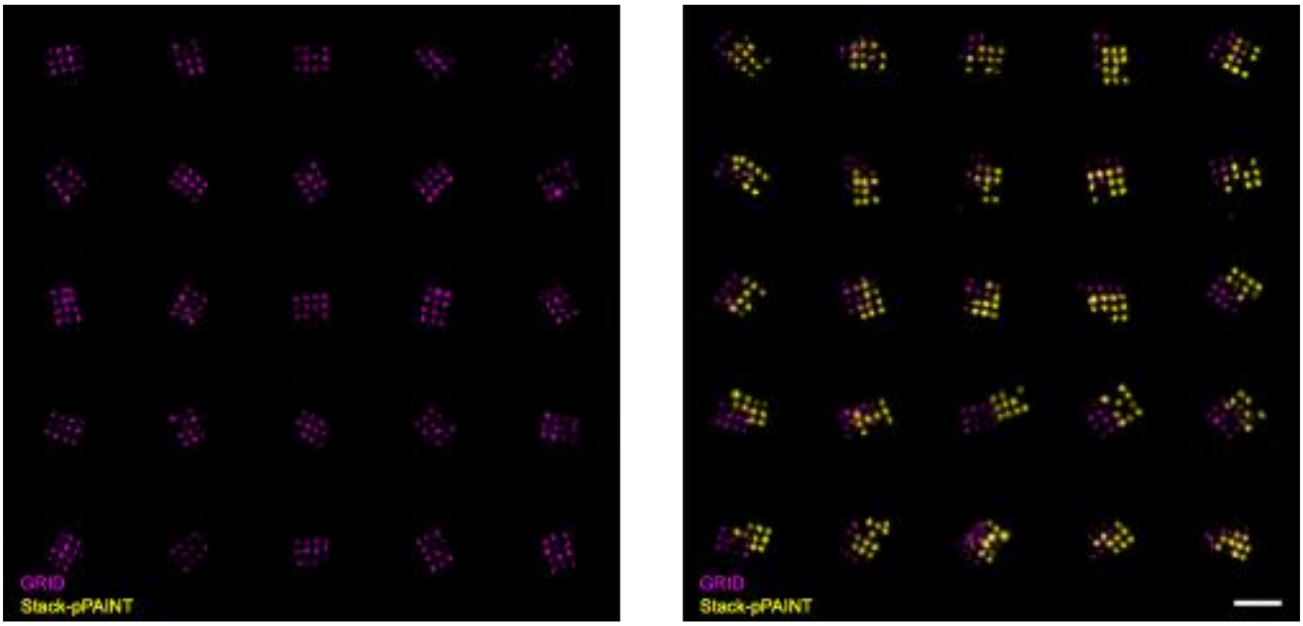


***Supplementary Figure 8: Example origami nanostructures for stack-pPAINT demonstration:*** *20 nm grid origami nanostructures with direct extensions (magenta) and stack-pPAINT probes (yellow). The origami nanostructures lacked (left) or had (right) the pST extensions. Note: Shifts in the two channels are due to chromatic aberrations observed under super-resolution.*

Supplementary Figure 9

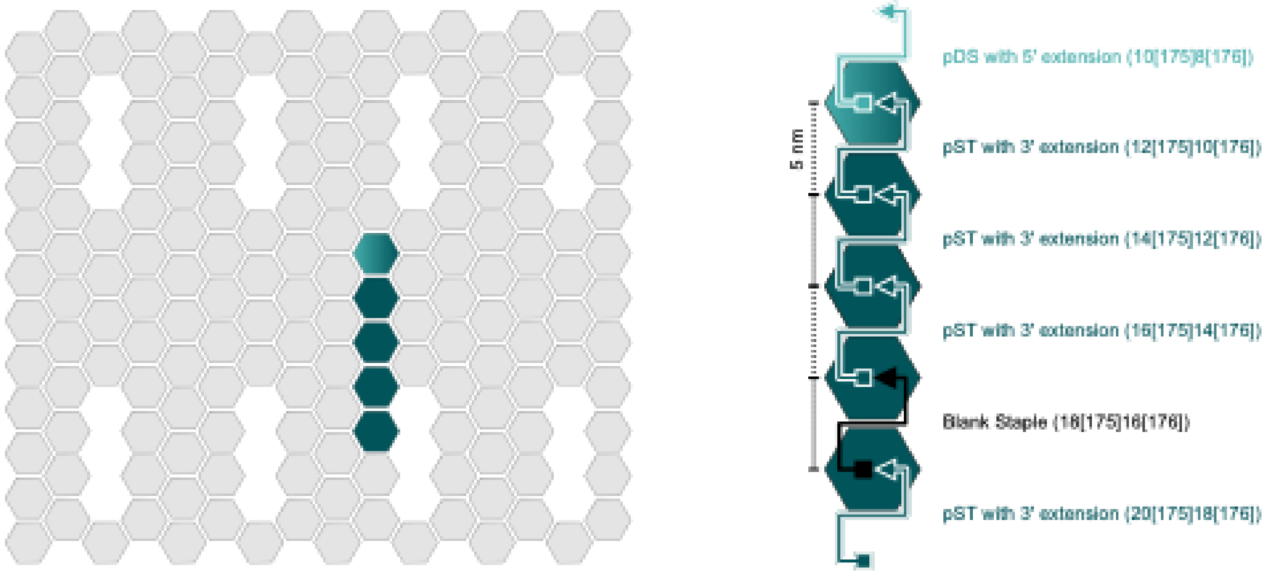


***Supplementary Figure 9: Stack-pPAINT efficiency assay staple layout:*** *Staples for assaying efficiency were places at the show site with a fixed position for pDS and varying postions for pST and 5 nm, 10 nm and 20 nm assay distances (D). The leash (L) and stem (S) lengths were varied as described.*

### Supplementary Figure 10

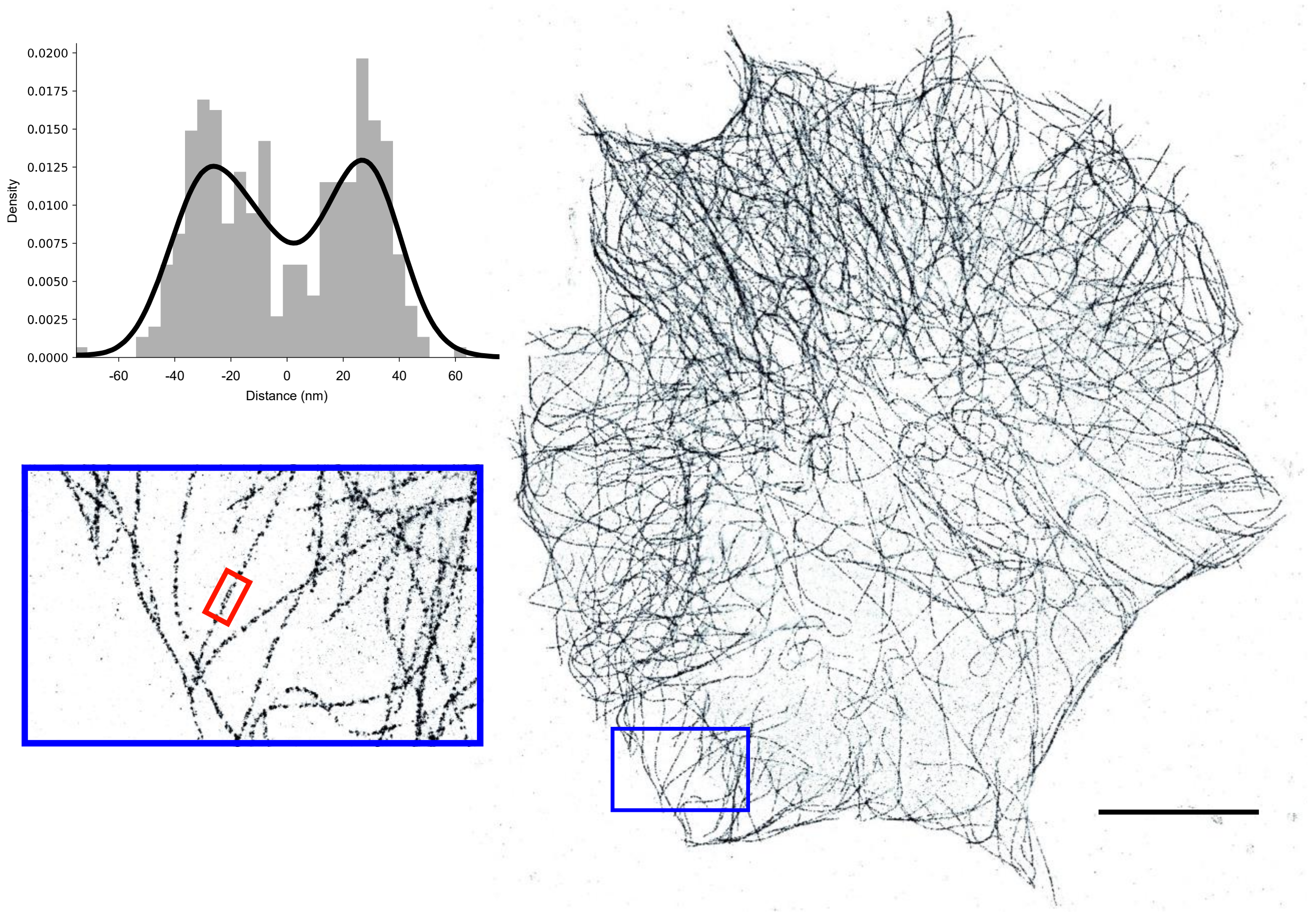


***Supplementary Figure 10: Microtubular network imaging with Stack-pPAINT:*** *Microtubules show an ~60 nm width. (Scale: 10 µm).*

### Supplementary Text

#### Average stacking of dye, terminal overhang and base-pairs in simulations

Average stacking percentages of the bases in simulation were computed based on a center of mass distance cutoff of 3Å and normal vector angle cutoff of 45°. Notably, Cy3B stacking directly with DC3, its attachment site, was observed to be a rare event across all systems and runs, indicating that such interactions are energetically or sterically unfavorable. Instead, we frequently observed displacement of DC3, with Cy3B preferentially stacking onto the adjacent first base pair of the duplex. This interaction appeared with varying degrees across systems and between independent replicas, suggesting that while this mode of stacking is possible, it is highly variable and sensitive to local structural context. The variability in stacking percentages further indicates that our simulations have captured only a small subset of rare events in the vast conformational landscape sampled in MD simulations. Interestingly, among the systems without Cy3B, system 2 exhibited the highest degree of stacking between DC3 and the first base pair, suggesting that in the absence of dye, DC3 more stably stacks onto the duplex. At the nick site, we observed consistently high base stacking across all systems, with system 3—lacking a physical nick—showing the highest stacking percentage. This suggests that the nick site retains intrinsic stacking stability, potentially contributing to duplex integrity despite backbone discontinuity (Supplementary Figure 3). One interesting thing to note here is, having Cy3B preserved Watson-Crick hydrogen bonds in our models suggesting that Cy3B's presence has impact on base pair stability.

#### Principal component analysis

To determine the major motion patterns in our all-atom models, we applied principal component analysis (PCA)[3]. This approach involves generating a covariance matrix from the simulation data and computing its eigenvectors. By projecting the trajectories along these eigenvectors, we can isolate and visually inspect the principal motions that dominate the system's dynamics. PCA was computed 6 atoms of every nucleotide in DNA (C1', C2, C3', C4, C5', N1) to capture the essential backbone and base dynamics of the DNA. The normalized population distributions along the top three principal components (PCs) for each system are shown in figure below. These projections reveal that the most dominant motions differ between systems, with distinct distribution patterns and populations observed along PC1–PC3.

### Supplementary Movies

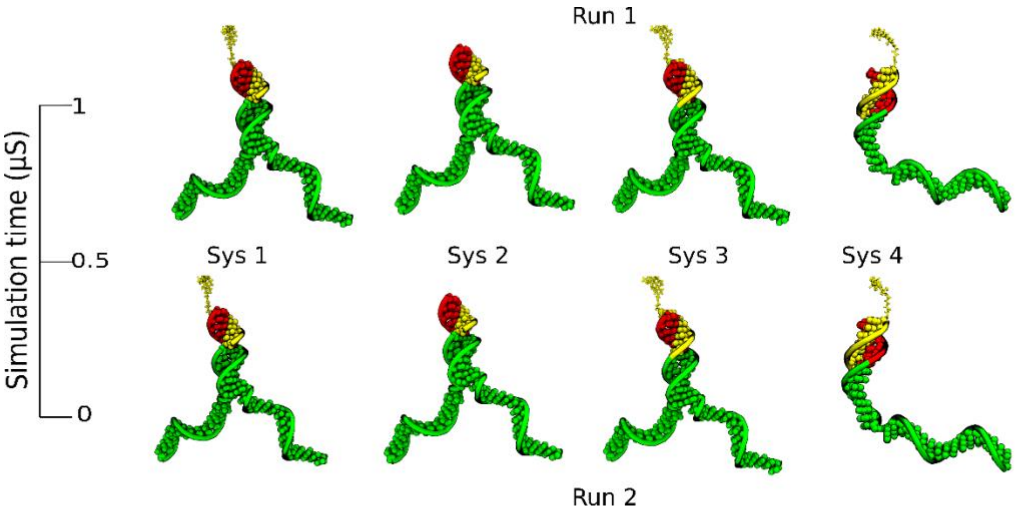


**Supplementary movie 1: Simulation trajectories of two MD runs of five systems mentioned in table 1:** This video illustrates 1 µs long equilibrium MD simulation trajectories of 5 systems under study with all-atom resolution. The atoms of 7 base imager and Cy3B molecules are shown using yellow spheres, docking strand which is complementary to imager is shown in red while the rest of docking strand is shown in green van der Waal's representation. Water and ions are not shown for the sake of clarity.

Online Link: Simulation Movie 1

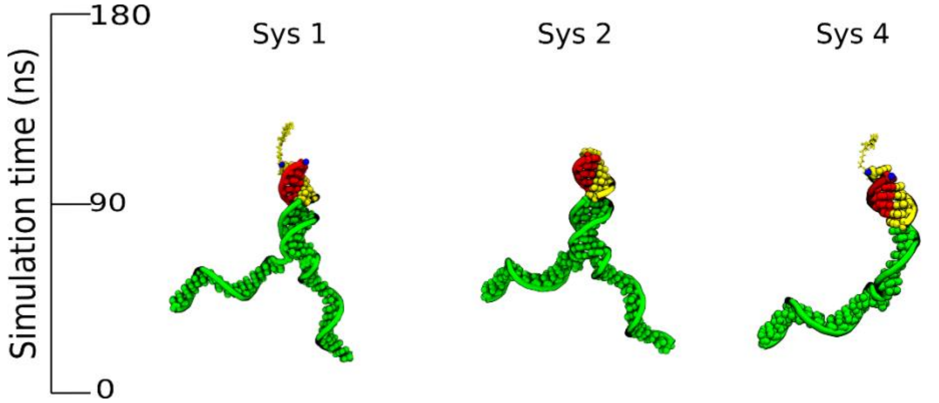


**Supplementary movie 2: Adaptive Steered MD trajectories:** This video illustrates 180 ns long adaptive steered MD simulation trajectories of system 1 2, and 4 with all-atom resolution, wherein imager is being pulled away from docking strand at the velocity of 0.5 Å/ns. Water and ions are not shown for the sake of clarity.

Online Link: Simulation Movie 2
